# Pipeliner: A Nextflow-based framework for the definition of sequencing data processing pipelines

**DOI:** 10.1101/476515

**Authors:** Anthony Federico, Tanya Karagiannis, Kritika Karri, Dileep Kishore, Yusuke Koga, Joshua D. Campbell, Stefano Monti

**Affiliations:** Bioinformatics Program, Boston University, 24 Cummington Mall, Boston, MA 02215; Division of Computational Biomedicine, Boston University School of Medicine, 75 East Newton Street, Boston, MA, 02118

## Abstract

The advent of high-throughput sequencing technologies has led to the need for flexible and user-friendly data pre-processing platforms. The Pipeliner framework provides an out-of-the-box solution for processing various types of sequencing data. It combines the Nextflow scripting language and Anaconda package manager to generate modular computational workflows. We have used Pipeliner to create several pipelines for sequencing data processing including bulk RNA-seq, single-cell RNA-seq (scRNA-seq), as well as Digital Gene Expression (DGE) data. This report highlights the design methodology behind Pipeliner which enables the development of highly flexible and reproducible pipelines that are easy to extend and maintain on multiple computing environments. We also provide a quick start user guide demonstrating how to setup and execute available pipelines with toy datasets.

## Introduction

High-throughput sequencing (HTS) technologies are vital to the study of genomics and related fields. Breakthroughs in cost efficiency have made it common for studies to obtain millions of raw sequencing reads. However, processing this data requires a series of computationally intensive tools which can be unintuitive to use, difficult to combine into stable workflows that can handle large number of samples, and challenging to maintain over long periods of time in different environments.

Pipeliner is a framework for the definition of sequencing data processing pipelines that aims to solve these issues. Pipelines developed within the framework are platform independent, fully reproducible, and inherit automated job parallelization and failure recovery. Their flexibility and modular architecture allows users to easily customize and modify processes based on their needs. Pipeliner also provides additional resources that allow developers to rapidly build and test their own pipelines in an efficient and scalable manner. Pipeliner is a complete and user-friendly solution to meet the demands of processing large amounts and various types of sequencing data.

## Materials and Methods

### Design and Features

Pipeliner is a suite of tools and methods for defining sequencing pipelines. It uses Nextflow, a portable, scalable, and parallelizable domain-specific language, to define data workflows (Di Tommaso *et al.*, 2017). Using Nextflow, each pipeline is modularized, consisting of a configuration file as well as a series of processes. These processes define the major steps in each pipeline, and can be written in Linux-executable scripting languages such as Bash, Python, Ruby, etc. Nextflow processes are connected through channels – asynchronous FIFO queues – which allow data to be passed between the different steps in each pipeline using a Dataflow programming model. Using this architecture, pipelines developed within the Pipeliner framework inherit multiple features that contribute to their flexibility, reproducibility, and extensibility.

### Pipeline Flexibility

Pipeliner enables flexible customization of pipeline options and parameters. Pipeliner currently offers three pipelines to demonstrate its applicability in processing different types of data, including bulk RNA-seq, single-cell RNA-seq (scRNA-seq), as well as Digital Gene Expression (DGE) data (Soumillon *et al.*, 2014). For the RNA-seq pipeline, sequencing reads are checked for quality with FastQC (Andrews, 2010), trimmed with TrimGalore (Andrews, 2012), mapped to a reference genome with either STAR (Dobin *et al*., 2012) or HISAT2 (Kim *et al*., 2015), and quantified with either StringTie (Pertea *et al*., 2015), HTSeq (Anders *et al*., 2015), or featureCounts (Liao *et al*., 2014). After alignment, mapping quality is checked with RSeQC (Wang *et al.*, 2012) and a comprehensive summary report of all processes is generated with MultiQC (Ewels *et al*., 2016). The scRNA-seq and DGE pipelines adopt a similar methodology, and the development of additional pipelines for microRNA-seq (miRNA-seq) and RNA-seq Variant Calling is currently underway.

### Parameter Configuration

All pipeline options and process parameters are set from a single configuration file (Fig. 2). Users have the option to select and skip various steps as well as customize parameters and allocate computing resources for specific processes. This flexibility gives rise to many different use cases. For example, a user may opt to provide a pre-indexed reference genome, or start the pipeline after the mapping step with saved alignment files, or output an ExpressionSet data structure with count and phenotypic data. Thus, each pipeline is multi-purpose and allows users to frequently tweak settings without adding complexity or sacrificing reproducibility.

**Fig 1.**
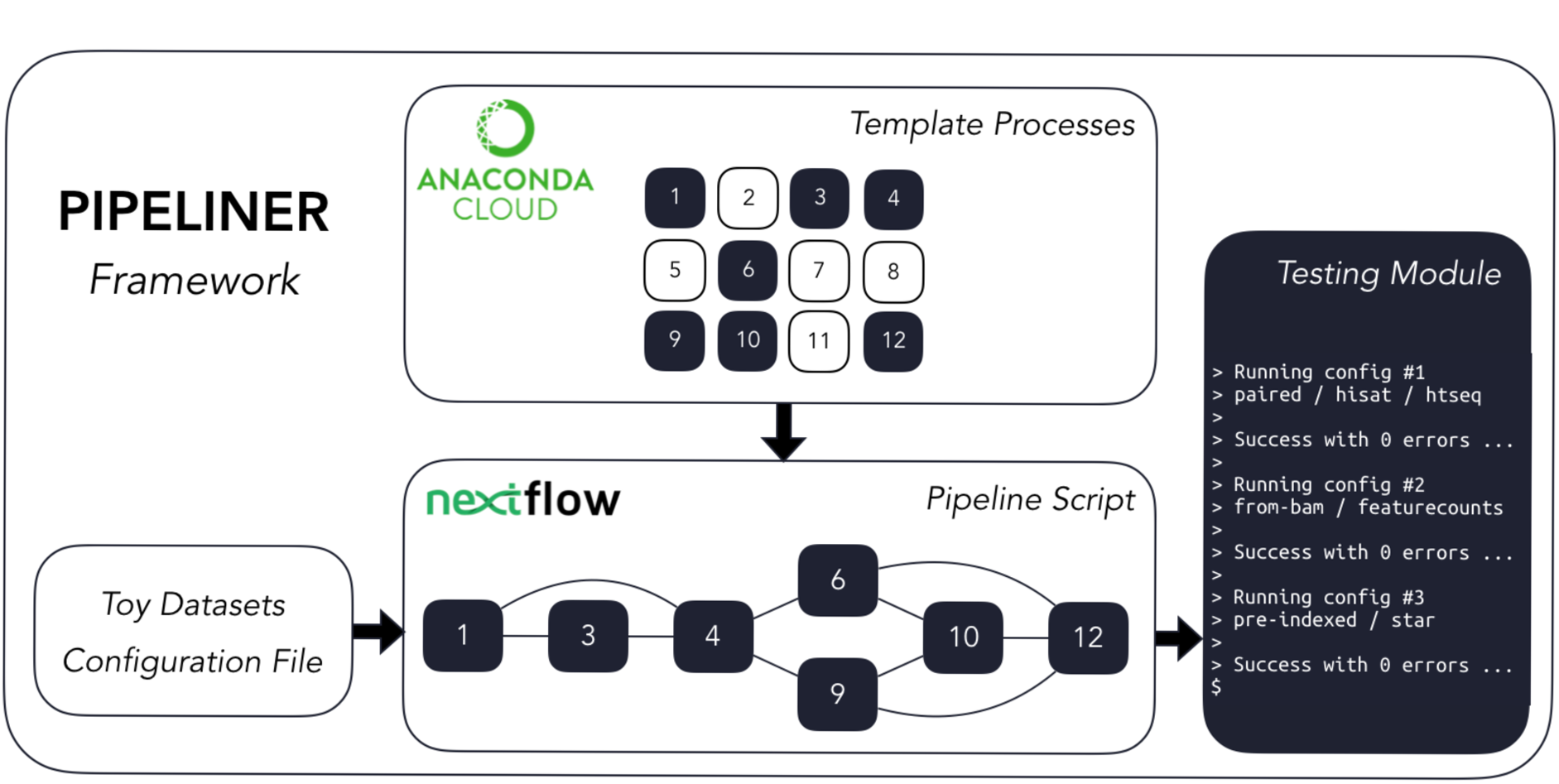
The Pipeliner framework employs reusable template processes strung together via Nextflow’s scripting language to create workflows in addition to developer tools such as toy datasets and testing modules.

**Fig 2.**
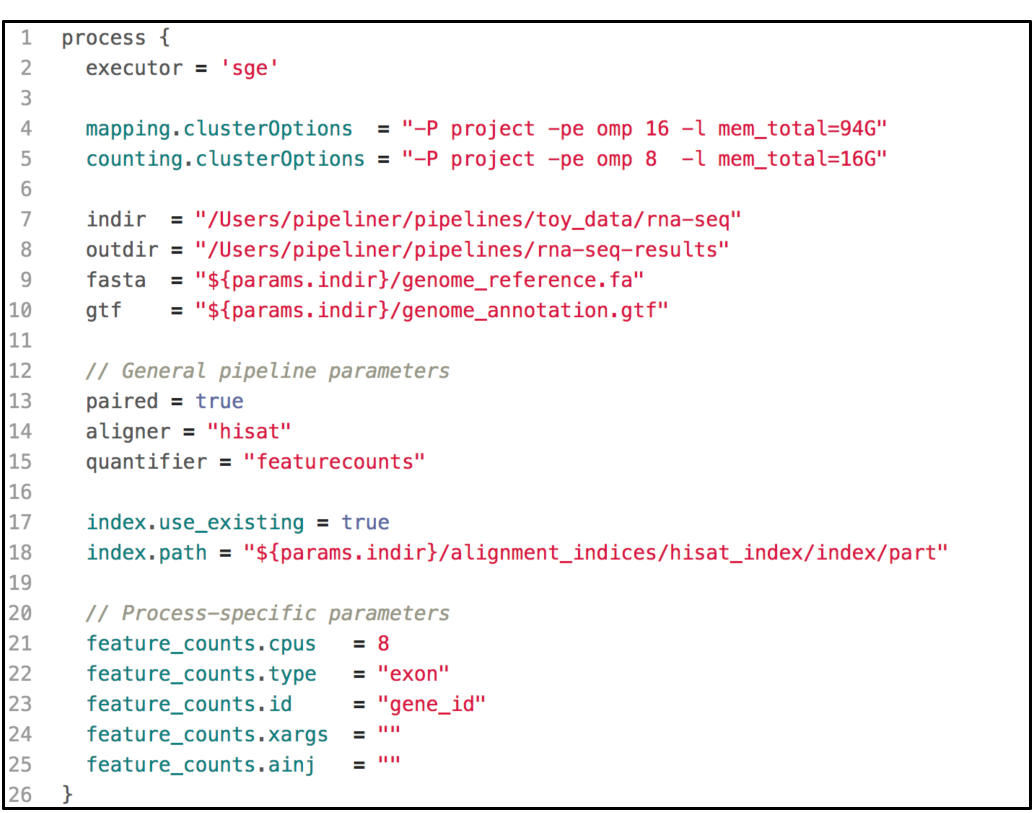
A shortened example of a configuration file, highlighting the key components. This configuration includes resource allocations for cluster executions, input and output paths to data, general pipeline parameters, as well as process-specific parameters.

The default configuration file defines variables for common parameters of third-party software tools used in each pipeline. These tools are wrapped into templates – one for each process – which are executed sequentially within the pipeline script. Because some software tools have hundreds of arguments, users have the option to insert code injections from the configuration file. These code injections can be used to pass uncommon keyword arguments or to append ad hoc processing steps (Fig. 3). These features provide unrestricted control over each step in the execution of a pipeline. Furthermore, since all modifications are made within the configuration file – which is copied with each run – the pipeline script is left intact, preserving the reproducibility of each run regardless of any execution-specific changes the user may make.

**Fig 3.**
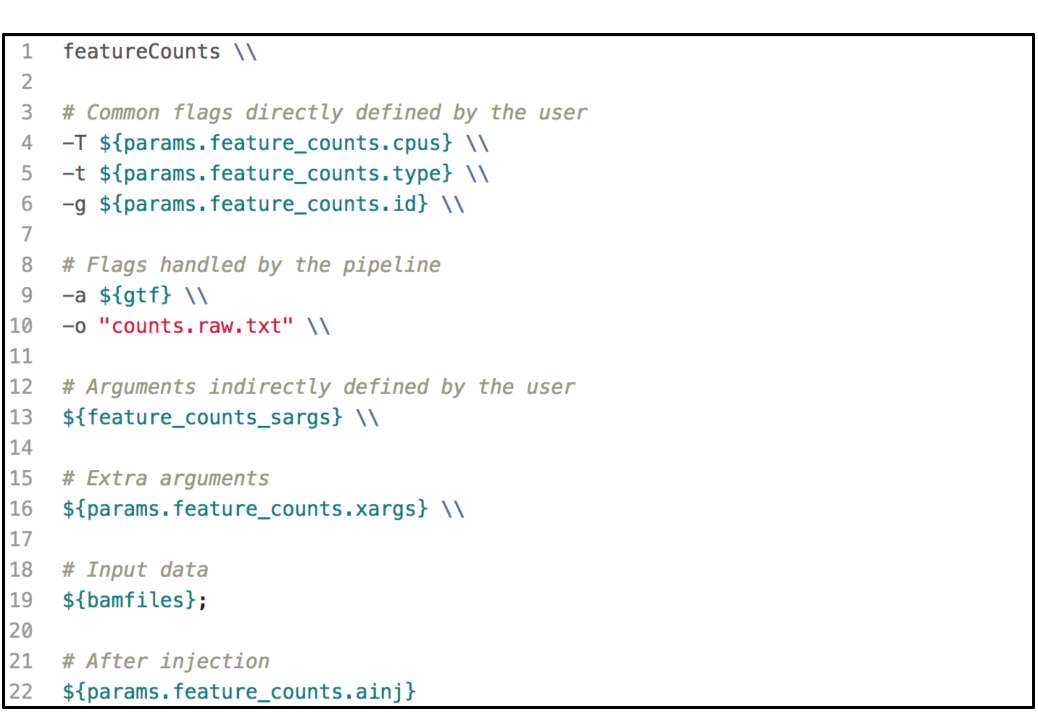
A code example of a template defined for the software tool featureCounts. The template wraps user-defined parameters, paths to data, as well as code injections into an executable bash script used in one of the pipeline steps.

### Workflow Reproducibility

Pipeliner is designed to create reproducible workflows. An abstraction layer between Nextflow and Pipeliner logic enables platform independence and seamless compatibility with high performance cloud computing executors such as Amazon Web Services (AWS). Pipeliner also uses Anaconda – a multi-platform package and environment manager – to manage all third-party software dependencies and handle pre-compilation of all required tools before a pipeline is executed (Continuum Analytics, 2016).

Pipeliner is bundled with a pre-packaged environment hosted on Anaconda Cloud which contains all software packages necessary to run any of the three pipelines available. This virtual environment ensures consistent versioning of all software tools used during each pipeline execution. Additionally, all file paths, pipeline options, and process parameters are recorded, time-stamped, and copied into a new configuration file with each run, ensuring pipelines are fully reproducible regardless of where and when they are executed.

### Extensibility

Pipeliner makes the development of Bioinformatics pipelines more efficient. The configuration file and processes that makeup each pipeline are inherited from shared blocks of code called template processes. For example, if a major update to an alignment tool requires modification to its template process, these changes propagate to all pipelines inheriting it. This property also minimizes the amount of code introduced as new pipelines are created, making them quicker to develop and easier to maintain. If a pipeline can inherit all of its processes with pre-defined templates, the user is only required to link these processes via Nextflow’s scripting language and create a basic configuration file.

## Rapid Development and Testing

Users can rapidly develop pipelines by using the toy datasets conveniently included with Pipeliner, enabling developers to test modifications made to their pipeline in minutes rather than hours. When testing, each execution covers only one configuration of parameters, meaning some processes may be skipped or partially executed depending on the configuration file. Therefore, to increase decision coverage, that is, the amount of tested reachable code, Pipeliner includes a custom testing module which automatically executes and logs a series of independent tests and configuration files (Fig. 5). With these tools, users can efficiently build, test, and maintain multiple sequencing pipelines.

**Fig 4.**
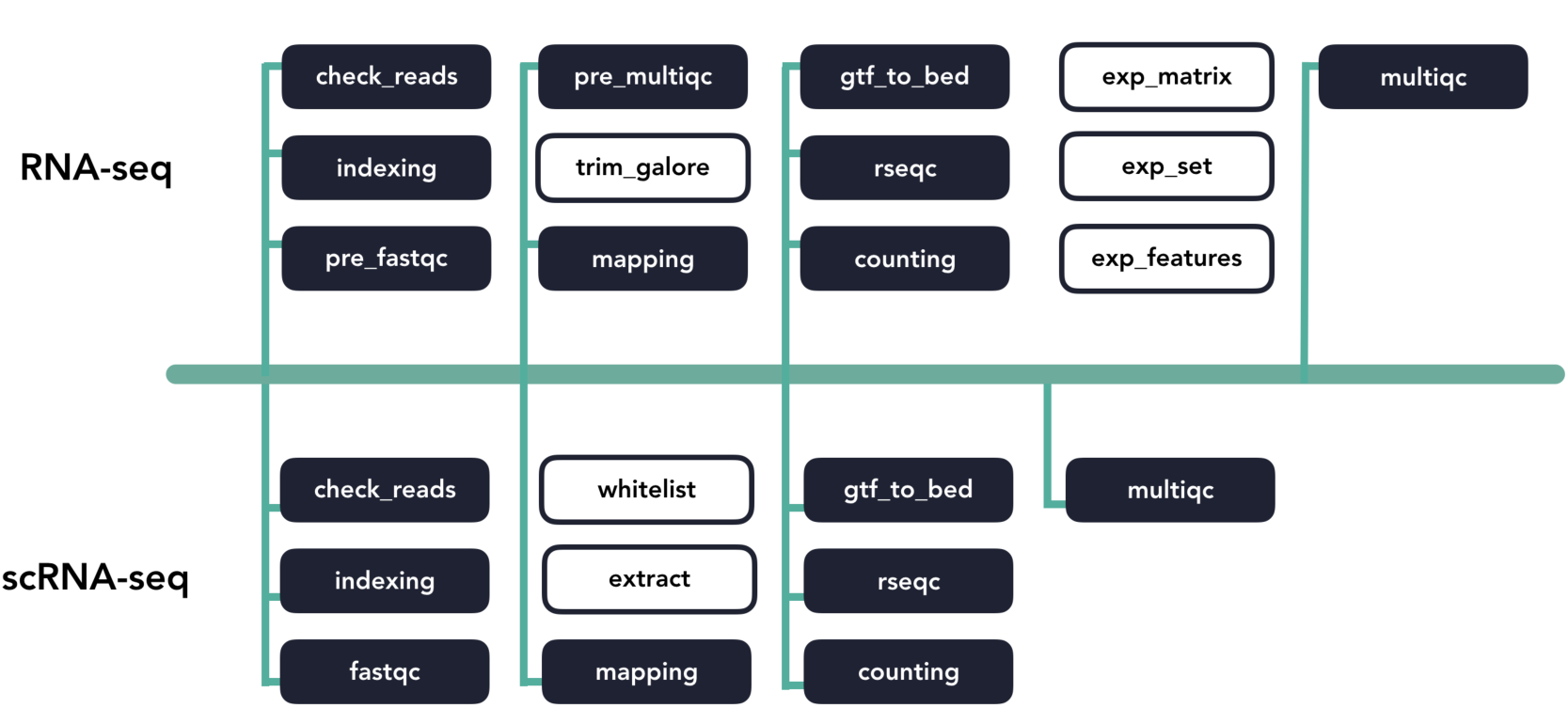
A diagram of template code sharing between the RNA-seq and scRNA-seq pipelines. Each block represents an individual workflow step. Shaded blocks share template code while unshaded blocks are unique.

**Fig 5.**
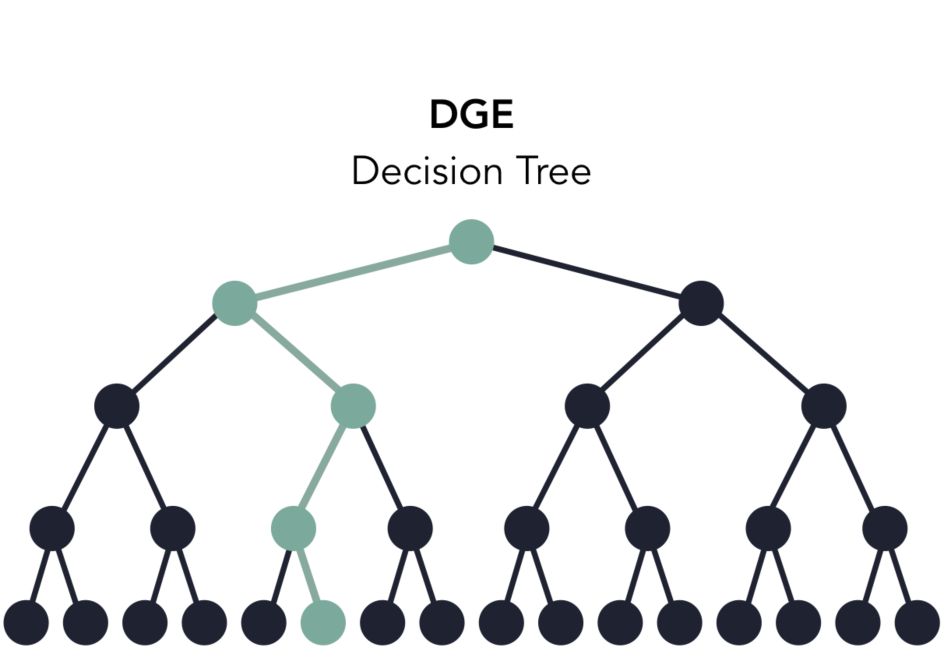
A decision tree of the DGE pipeline based on available options in the configuration file. When testing, one execution of the pipeline will test one path in the decision tree (highlighted in green). The testing module automates the execution and logging of all possible paths.

## Usage Guide

In addition to <u>comprehensive documentation</u> of the framework and to demonstrate its ease of use, we provide a tutorial for processing the toy datasets available for each pipeline.

### Processing Toy Datasets

The Pipeliner framework requires Nextflow and Anaconda. Nextflow requires <u>Java 8 (or higher)</u> to be installed and can be used on Linux and OS X machines. Third-party software tools will be installed and managed through an Anaconda virtual environment. Once the pre-requisites are installed, the repository can be cloned from GitHub to any location through the following command:

~~~
$ git clone https://github.com/montilab/pipeliner
~~~

The next step is to clone and activate the virtual environment. The easiest method is to recreate the environment through the yml files provided in the repository. There is a single yml file for both Linux and OS X operating systems, containing all dependencies for all available pipelines.

~~~
$ conda env create-f pipeliner/envs/linux_env.yml # Linux
$ conda env create-f pipeliner/envs/osx_env.yml # OS X
$ source activate pipeliner
~~~

Pipeliner requires configuration of paths to input data such as fastq reads, bam alignments, references files, etc. When cloning Pipeliner to a new machine, all paths must be reconfigured. This process can be automated by running a script which will reconfigure any paths to the same directory of your clone.

~~~
$ python pipeliner/scripts/paths.py
~~~

The final step is to download a Nextflow executable package in the same directory as the available pipelines.

~~~
$ cd pipeliner/pipelines
$ curl-s https://get.nextflow.io|bash
~~~

With the setup complete, any of the available pipelines can be executed with their respective toy datasets with the following commands.

~~~
$ ./nextflow rnaseq.nf-c rnaseq.config
$ ./nextflow scrnaseq.nf-c scrnaseq.config
$ ./nextflow dge.nf-c dge.config
~~~

### Proof of Concept

To display the applicability of Pipeliner to real-world datasets, we reprocessed 48 RNA-seq paired read files for the Lymphoid Neoplasm Diffuse Large B-Cell Lymphoma (DLBC) cohort from The Cancer Genome Atlas (TCGA). For each cohort, the TCGA uses a standardized pipeline where reads are mapped to a reference genome with STAR and quantified by HTSeq. While the TCGA provides open access to the count matrix, some researchers have opted to use alignment and quantification algorithms specific to their research interests (Mumtahena *et al*., 2015). For this reason, the TCGA also provides raw sequencing data, however its large size requires parallelization on a high-performance computing platform. We argue Pipeliner is a suitable choice for users looking for alternative reprocessing of TCGA datasets with minimal pipeline development.

Pipeliner makes alternative processing of TCGA and other publicly available data straightforward. In processing raw RNA-seq data for DLBC, paired fastq reads were downloaded from the Genomic Data Commons (GDC) Data Portal. For each sample, Pipeliner requires an absolute file path to reads. After specifying this information, Pipeliner was able to successfully process all data with HISAT2, featureCounts, and the remaining settings left to default (Fig. 6). The flexibility provided by Pipeliner is ideal for users experimenting with different tools and parameters. For example, because Pipeline is capable of taking aligned bam files as input and skipping preceding steps, we were able to rapidly try all three quantification options without re-running unrelated processes. This level of control is critical for downstream analysis of the processed data. To help researchers extend this example to other datasets, we provide the scripts used to obtain and organize TCGA data from the GDG as well as the configuration file used by Pipeliner to process the data in the supplementary information.

**Fig 6.**
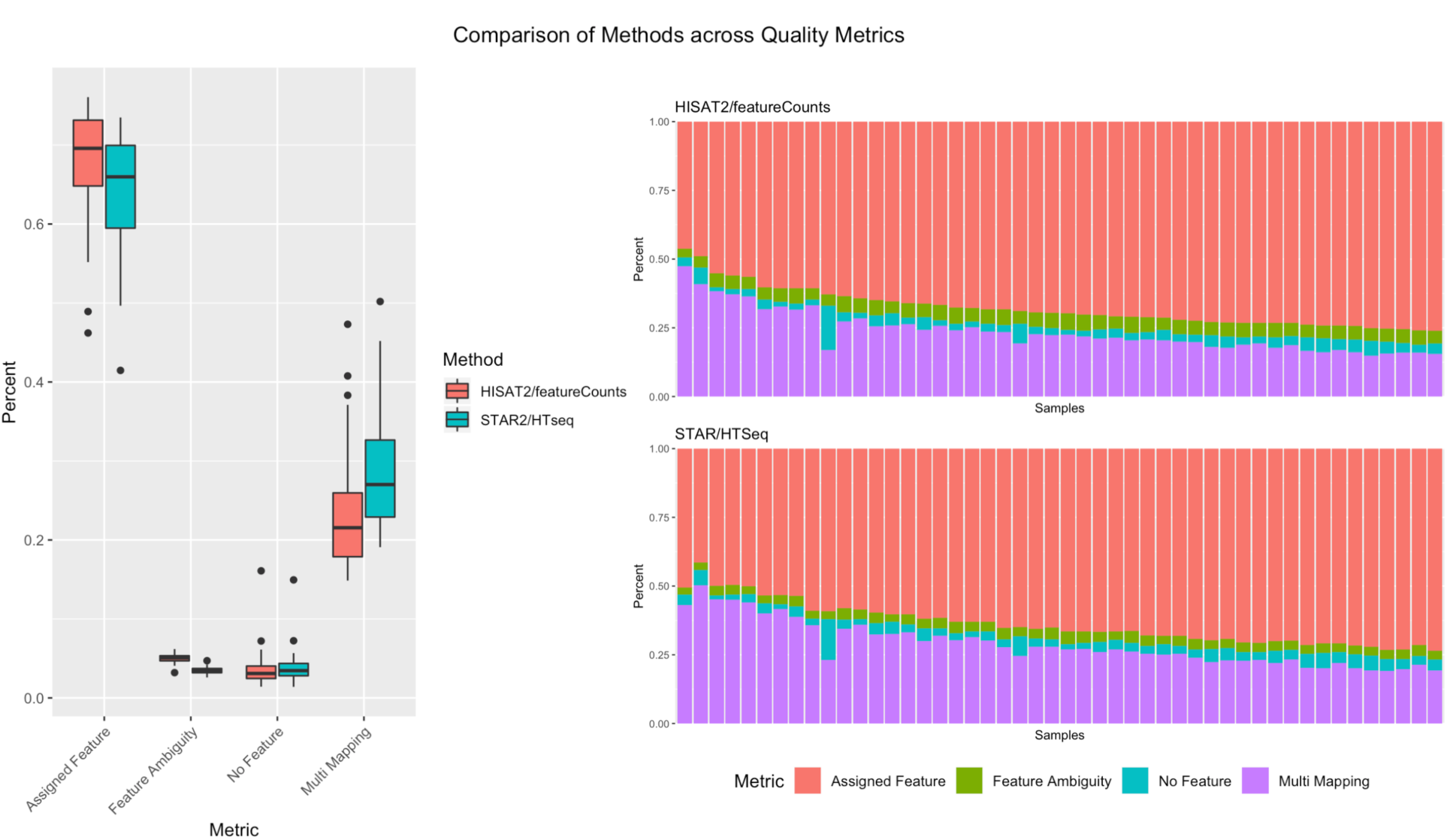
A summary of counting and mapping quality metrics when processing with Pipeliner using HISAT2 and featureCounts compared with processed counts from TCGA-DLBC using STAR and HTSeq.

## Conclusions

Together with Nextflow and Anaconda, Pipeliner enables users to process large and complex sequencing datasets with pipelines that are customizable, reproducible, and extensible. The framework provides a set of user-friendly tools for rapidly developing and testing new pipelines for various types of sequencing data which will inherit valuable design features of existing pipelines. We apply the RNA-seq pipeline to real-word data by processing raw sequencing reads from the DLBC cohort provided by the TCGA and provide supplementary files which can be used to repeat the analysis or serve as a template for applying Pipeliner to other publicly available datasets.

## Availability and Future Directions

Pipeliner is implemented in Nextflow, Python, R, and Bash and released under an MIT license. It is publicly available at https://github.com/montilab/pipeliner and supports Linux and OS X operating systems. Comprehensive documentation is generated with Sphinx and hosted by Read the Docs at https://pipeliner.readthedocs.io/. We will continue to develop the Pipeliner framework as the Nextflow programming language matures and we plan to provide additional pipelines for other types of sequencing data and analysis workflows in the future.

## Supporting information

## Acknowledgements

The authors would like to thank P. Di Tommaso for his assistance with Nextflow-related inquiries and A. Gower for his advice for improving and testing Pipeliner.

