## Supplementary material for "Pipeliner: A Nextflow-based framework for the definition of sequencing data processing pipelines": 02_naming.py

```
#!/usr/bin/env python
import os

path = os.environ['WORK']

samples = {}
with open(path+'/SAMPLES.txt', 'r') as infile:
    for i, line in enumerate(infile):
        if i == 0:
            continue
        line = line.split()
        samples[line[0]] = line[3]

with open(path+'/DLBC_fq.csv', 'w') as outfile:
    outfile.write('Sample_Name,Read1,Read2\n')
    for l1 in os.listdir(path+'/DLBC'):
        for l2 in os.listdir(path+'/DLBC/'+l1):
            if l2.endswith('analysis.xml'):
                with open(os.path.join(path+'/DLBC/'+l1, l2), 'r') as infile:
                    for line in infile:
                        line = line.strip('\n').strip('\t')
                        if line.startswith('<DESCRIPTION>'):
                            fid = line.split(' ')[-1].split('<')[0]
                            sid = samples[fid]
            if l2.endswith('1.fastq'):
                r1 = path+'/DLBC/'+l1+'/'+l2
            if l2.endswith('2.fastq'):
                r2 = path+'/DLBC/'+l1+'/'+l2
        outfile.write('{0},{1},{2}\n'.format(sid, r1, r2))
```
