## Supplementary material for "Pipeliner: A Nextflow-based framework for the definition of sequencing data processing pipelines": README.txt

This file provides information regarding the scripts used to download and 
organize TCGA data from Genomics Data Commons (GDC) in the context of the DLBC cohort.

STEP #1 - 01_download.sh

This script uses the gdc-client tool to download raw data from the GDC. It uses an environment variable called $WORK, which should be set to your working directory. Also required is a MANIFEST.txt file (a comma-separated list of UUIDs) as well as a TOKEN.txt file (an authentication token). Both files can be obtained from the GDC data portal after logging in. Please refer to their documentation at https://docs.gdc.cancer.gov/API/Users_Guide for more information. Note, make sure the DLBC directory exists in $WORK before running the script.

Before moving to the next step, decompress the .tar.gz files within the sub directories of $WORK/DLBC. Because this process may take a long time, you may want to parallelize it, which is why a script is not included. Make sure to keep decompressed files in the same directory as their compressed version.

For example:
tar -zxf path/to/dir/filename.tar.gz --directory path/to/dir

STEP #2 - 02_naming.py

This script is used to name each sample and create a csv file linking sample names to absolute file paths to each read. This file is used by Pipeliner to load in read files and reference the samples they belong to. It requires a SAMPLES.txt file which contains sample information regarding the TCGA cohort of interest and can be obtained from the GDC data portal. Please refer to their documentation at https://docs.gdc.cancer.gov/API/Users_Guide for more information.
